# Optimising source identification from marmoset vocalisations with hierarchical machine learning classifiers

**DOI:** 10.1101/2022.11.19.517179

**Authors:** Nikhil Phaniraj, Kaja Wierucka, Yvonne Zürcher, Judith M. Burkart

## Abstract

Marmosets, with their highly social nature and complex vocal communication system, are important models for comparative studies of vocal communication and, eventually, language evolution. However, our knowledge about marmoset vocalisations predominantly originates from playback studies or vocal interactions between dyads, and there is a need to move towards studying group-level communication dynamics. Efficient source identification from marmoset vocalisations is essential for this challenge, and machine learning algorithms (MLAs) can aid it. Here we built a pipeline capable of plentiful feature extraction, meaningful feature selection, and supervised classification of vocalisations of up to 18 marmosets. We optimised the classifier by building a hierarchical MLA that first learned to determine the sex of the source, narrowed down the possible source individuals based on their sex, and then determined the source identity. We were able to correctly identify the source individual with high precisions (87.21% – 94.42%, depending on call type, and up to 97.79% after the removal of twins from the dataset). We also examine the robustness of identification across varying sample sizes. Our pipeline is a promising tool not only for source identification from marmoset vocalisations but also for analysing vocalisations and tracking vocal learning trajectories of other species.

## INTRODUCTION

Comparative studies of primate communication are crucial for understanding the evolutionary origins of human speech and language as they allow for identifying the features of speech and language that we share with non-human primates [1–5]. A long tradition of comparative studies has found that some primates have the anatomical capabilities of producing human-like speech [6,7], can combine call types into syntactically ordered sequences [8], have basic phono-articulatory coupling potential [9–11] and show some vocal plasticity [12–14]. The vocal communication of callitrichid monkeys appears particularly rich among non-human primates. For instance, common marmosets (*Callithrix jacchus*) have been shown to possess superior control and flexibility in their calls. Among others, they can actively interrupt ongoing phee calls [15,16], make long-term changes to call frequency in response to noise [15], and converge in vocal space to a social partner [12]. As immatures, they go through a babbling phase [17], and social inputs and contingent vocal feedback from caregivers contribute to fully developing their repertoire [18–20]. Marmosets have a broad vocal repertoire and produce a large variety of compound calls formed by using one or more simple call types uttered in a sequence [21]. Trills are the most common call type and are used as within-group close contact signals; phees are louder long-distance contact calls, and food calls are elicited by individuals when they see food, during feeding, and to initiate food sharing [22].

Marmosets are not very closely related to humans. However, they are similar to us with regard to how they raise offspring: like humans, they are cooperative breeders, i.e., they have a reproductive system in which group members other than the parents significantly contribute to raising offspring. Such a system requires high levels of collective action and group coordination in turn requiring exceptional communicative capabilities. This convergence may thus have played a role in language evolution [23–26], making marmosets particularly relevant primate models. However, most studies on marmoset vocal communication are performed under highly controlled conditions, often focusing on a single individual only or physically separated dyads that can interact only vocally with each other. Only rarely are group-level communication dynamics taken into account [27]. Given the highly social nature of marmosets, the next frontier in studying marmoset vocal communication is to do so under more naturalistic conditions in social groups, where they display their communication skills in entirety.

A major bottleneck for studying group-level communication in marmosets and other animals lies in the ability of the researcher to accurately determine and label the identity of the caller (source identity) when recording the vocalisations from large groups. Such a task being laborious, has attracted the attention of signal processing experts and data scientists for its automation [28]. This has led to the development of a plethora of machine learning (ML) techniques for source identification, each competing for higher accuracy [29–32]. The automation might be trivial in some species, such as the songs of male zebra finches, where individual signatures are visible on the spectrograms of songs and can be told apart by a trained human eye [33]. The same task can be challenging when spectrograms of vocalisations from different individuals look similar to the human eye, such as in the case of many primates [34–37], including marmosets [21,38].

For automating acoustic data processing for marmosets, recent studies have trained machine learning algorithms (MLAs) to classify marmoset calls into different call types [39,40]. Although this is an essential step towards automation, the task itself is computationally not very demanding, as the call types look very different on the spectrogram and are easily distinguishable manually. On the other hand, calls of the same type produced by different individuals look very similar on a spectrogram. Therefore, the major challenge lies in automating the classification of calls based on source identity.

The goal of our study is to automatically determine source identity from marmoset calls with high accuracy. To do so, we address two issues that would potentially increase the accuracy. First, traditional approaches for animal vocalisation analysis may not capture enough variation of the calls, and second, hierarchical ML classifiers that take advantage of sex differences in calls may outperform non-hierarchical ones.

First, the traditional approach for the acoustic analysis of animal vocalisation analysis involves applying short-time Fourier transforms on the acoustic waveform to bring it to the frequency domain [41]. A handful of predetermined acoustic features are then extracted [42,43]. These features are typically chosen because we understand how modifications of the sound affect these feature values, making them easy to interpret [44–46]. A drawback of this approach is that not all animal vocalisations show maximum variability along these feature values. The small feature space and reduced class separability of calls due to limited features prevent ML approaches from achieving their maximum potential for obtaining high classification accuracies. In this paper, to overcome this shortcoming, we implement time series analysis directly on the acoustic waveform for plentiful feature extraction and tree-based classifiers for meaningful feature selection. This allows us to address the first issue by providing much more detailed representations of the calls than the traditional approach.

Second, we hypothesise that using a hierarchical ML classifier will improve classification accuracies by breaking up the classification problem into a hierarchy of smaller ones. Callitrichid vocalisations contain information about the sex of the caller [47–49], which can potentially assist MLAs with efficient source identification. We, therefore, develop a hierarchical ML classifier that first determines the sex of the source, thus narrowing down the possible source individuals, and then determines the source identity. We apply it to trills, phees, and food calls of 20 marmoset individuals and compare its performance to a non-hierarchical approach. We then analyse the acoustic features used by the hierarchical classifier for the various steps in the hierarchy and finally compare the precisions and recalls of the non-hierarchical and hierarchical classifiers at different sample sizes (i.e. number of calls per individual).

## METHODS

### Experimental Subjects

This study used marmoset vocalisations collected by Zürcher et al. [12]. The data contained vocalisations from 20 adult common marmosets, 10 males and 10 females. They were housed in 2.4m x 1.5m x 0.8m enclosures along with at least one other individual, with each group having access to a personal outdoor enclosure of the same dimension and a common experimental room. Lighting was regulated to maintain a 12/12 hour day/night cycle. Animals were fed a predetermined amount of vitamin-enriched mush in the morning, vegetables and fruits during noon, and one of either gum, boiled egg, cottage cheese, or insects after noon. Ad libitum access to water was always provided. All experiments were approved by Zürich’s cantonal veterinary office (licence ZH223/16).

### Vocalisation recordings and segmentation

Individuals were recorded in their home enclosures on multiple days spread over two to three weeks. Each recording session was about 30 minutes long. A condenser Microphone (CM16/CMPA, Avisoft Bioacoustics, Germany) connected to Avisoft UltraSoundGate 116H (Avisoft Bioacoustics, Germany) was used for recordings and calls were labelled in real time using Avisoft Recorder (Avisoft Bioacoustics, Germany). Calls were visualised and segmented using Avisoft Pro (Avisoft Bioacoustics, Germany). See [12,50] for detailed information about the recording procedure and processing.

### Datasets and feature extraction

As the original dataset was imbalanced (number of calls per call type per individual was highly variable), a combination of majority class random undersampling and Synthetic Minority Oversampling TEchnique (SMOTE) [51,52] was used to create the following 18 datasets (one original and five generated, for each of the three call types). SMOTE is a data augmentation technique that synthesises new data for minority classes to make them equal to the majority class. X = T for trills, P for phees, F for food calls in the name of the dataset.

1. Original-X: The original dataset after HCTSA processing.
2. Imbalanced-X: Marmosets with <25 calls per call type were removed from the Original-X datasets. No undersampling was done.
3. Balanced-X: SMOTE was applied on Imbalanced-X to obtain balanced datasets.
4. Balanced197-X: From the Imbalanced-X dataset, classes with >197 calls were undersampled to 197 calls. SMOTE was applied to this.
5. Balanced99-X: From the Imbalanced-X dataset, classes with >99 calls were undersampled to 99 calls. SMOTE was applied to this.
6. Balanced50-X: From the Imbalanced-X dataset, classes with >50 calls were undersampled to 50 calls. SMOTE was applied to this.

The smaller Balanced197-X, Balanced99-X, and Balanced50-X datasets were used to test the capability of the ML approach to classify calls in limited sample size scenarios.

Recent advances in time series analyses allow for performing multiple operations that provide meaningful information about the nature of the time series [53]. This can be applied to acoustic data by viewing the acoustic waveform as a time series of pressure points. We used the Highly Comparative Time Series Analysis (HCTSA) [54] toolbox on MATLAB [55] for feature extraction from marmoset calls. Features common across calls of all individuals for a given call type were used for further analyses. We visualised the trill, phee, and food call datasets using t-distributed Stochastic Neighbour Embedding (t-SNE) [56].

We trained individual multiclass Adaptive Boosting algorithms with decision trees as weak learners and 10-fold cross-validation for determining the important features to use for classification and for performing the classification itself. This method uses multiple weak learners to create a strong learner [57] and is considered better than random forest classifiers due to its higher accuracy and lower susceptibility to overfitting in certain cases [58,59]. We first trained AdaBoost on Imbalanced-X, Balanced-X, Balanced197-X, and Balanced99-X datasets to classify calls based on source identity – as a direct or “non-hierarchical approach” (in contrast to the hierarchical approach that was used later) to determine source identity from calls (using MATLAB’s ‘fitcensemble’, ‘AdaboostM1’, and ‘AdaboostM2’ functions). We chose the number of trees and learning rate based on the observations of the classification loss function.

Although classification accuracy is the most widely used metric to assess ML models, it is not a good representation of the model’s performance on class-imbalanced datasets. This is because the classifier can get away with a high accuracy score by simply predicting most data points as belonging to the majority class. In such cases, the Receiver Operating Characteristic (ROC) curve can be used to evaluate the classifier’s performance. The ROC curve visualises how the true positive rate changes as a function of the false positive rate at various threshold values. The area under this ROC curve, simply ‘Area Under Curve’ (AUC), can be a useful tool for examining classifier performance along with accuracy. It is important to note here that ROC-AUCs represent how well the MLA can separate a positive class from the negative classes and do not represent how well the MLA would perform classification when actually handed the task. Therefore, for assessing the performance of AdaBoost on Imbalanced-X datasets, ROC-AUC was calculated in a one vs rest (one positive, rest negative classes) setting for each class in the dataset, along with the accuracies. Even while assessing the performance of MLAs on balanced datasets, accuracy does not represent how variable the predictions for each of the classes (marmoset individuals) are. Therefore, for the rest of the classifiers, precisions and recalls for each class were calculated, and the summary of these values was used to assess their performance.

For inspecting if individual variability of calls within marmoset groups can be explained by variation in sex and whether MLAs could exploit this to perform better, individual AdaBoosts were trained on Imbalanced-X and Balanced-X datasets to classify calls based on sex, and then the source individual. Sex was chosen as a cue because previous studies in Callitrichids have shown that they can discriminate calls based on the sex of the source [47–49], and in our case, the total number of classes could be split into half based on the sex of the individual (10 males and 10 females out of 20 individuals). Later, each dataset was divided into 2 sub-datasets based on the ‘true’ sex of the source individual, and separate AdaBoosts were trained on each of them. This was the hierarchical approach to determine first the sex, and then the source identity from calls (figure 1). To assess the performance of the classifiers for the hierarchical classification approach, precisions and recalls for each individual were calculated for all classifiers as:

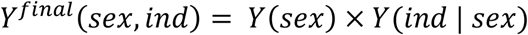

**Figure 1.**
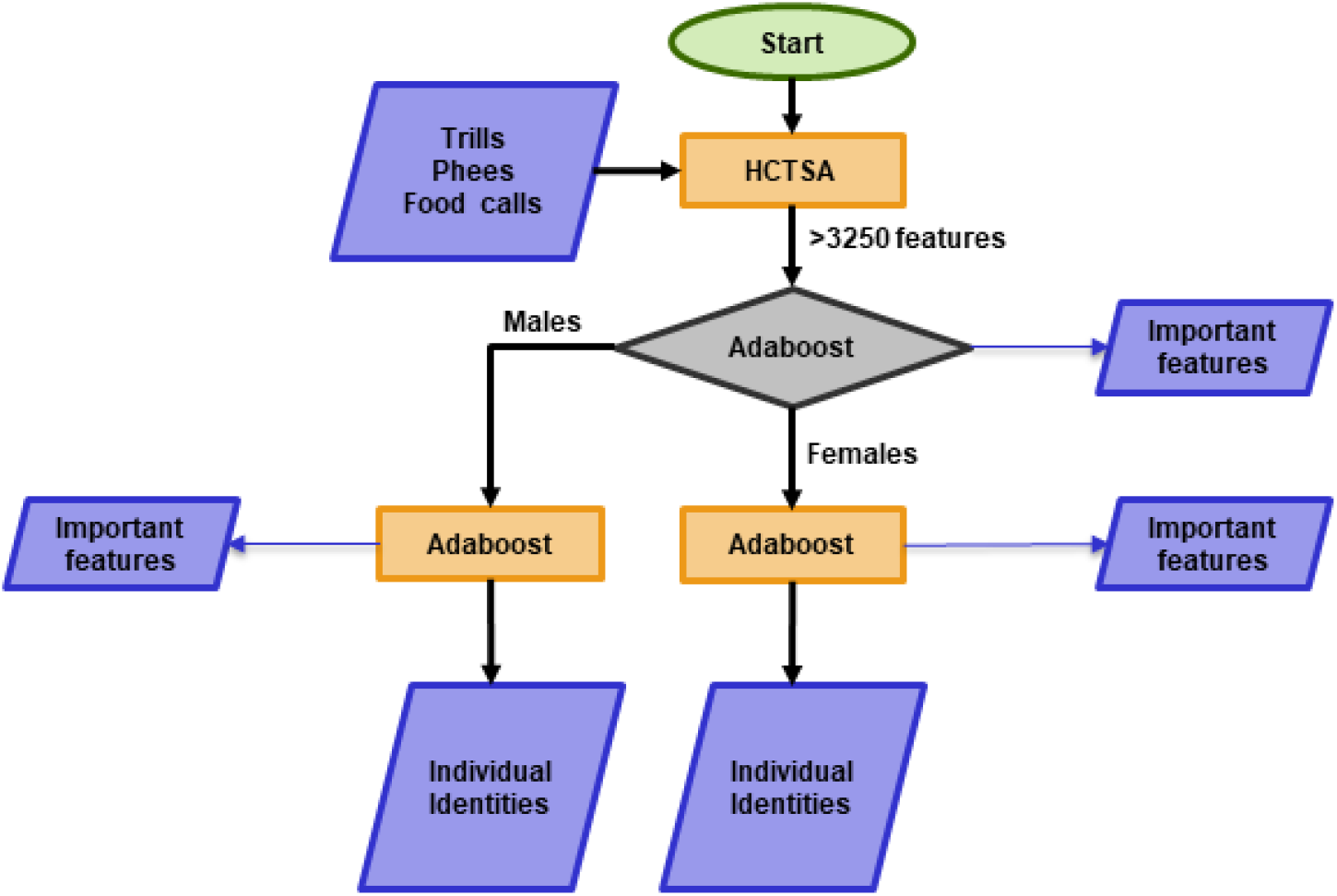
The hierarchical classification approach. Features from trills, phees, and food calls were extracted using HCTSA. These features were used to train AdaBoost to first classify calls based on sex, and then individual identities. The oval denotes start, parallelograms inputs/outputs, rectangles processes, and the diamond a decision.

*Y* is the precision or recall, and *ind* is the individual identity of the marmoset. For example, the final precision for a female individual would be the precision for determining the sex as a female, multiplied by the precision for determining the source individual among females.

Because of the difference in how the non-hierarchical and hierarchical approaches utilise the dataset for training, the usual pairwise t-test for mean precisions and recalls across the 10 folds could not be performed. Instead, we performed Wilcoxon signed-rank test to compare precision and recall scores for every class between the two approaches because the classes, i.e., the individuals for both the approaches were the same.

### Feature importance scoring

Along with classifying calls, each fold of the AdaBoost also provides predictor importance scores for each feature, which represents how important that feature was for AdaBoost in the classification task. First, we checked how (non-hierarchical) AdaBoost performed the feature selection task. For this, we ranked features by predictor importance scores given by the most accurate of the 10 models (run on 10 different folds) in the AdaBoost for each dataset. Then, for the 3 Balanced-X datasets, we used the top 20 features to visualise t-SNE clusters. t-SNE plots were generated using MATLAB’s ‘tsne’ function with the Barnes-Hut algorithm keeping the Barnes-Hut trade-off parameter at 0.5 to increase processing speed for large datasets. Exaggeration was set to 4, perplexity to n/100, and learning rate to n/12 (where n is the total number of calls for that call type, not to be confused with sample size per class) as these values are shown to provide robust results, which are comparable to UMAP [60], when datasets are large [61]. We compared these to t-SNE plots of the corresponding datasets, generated using 20 random features. For a more quantitative comparison, we selected 20 random features from each of the three datasets 100 times, plotted the histograms of the mean silhouette scores, fit Gaussians to these distributions, and calculated the probability of getting a mean silhouette score greater than that of the top 20 features by chance.

For the hierarchical classifiers, we analysed the features used by the various levels in the hierarchy when implemented on the 3 Balanced-X datasets. We checked what proportion of the total features utilised by the hierarchical classifier: (1) was used for determining the sex only, (2) was used for determining source identity among females only, (3) was used for determining source identity among males only, (4) were common between determining sex and individual identity among females, (5) were common between determining sex and individual identity among males, (6) were common between determining individual identities among females and males, and (7) were common between determining sex, individual identities among females, and individual identity among males.

## RESULTS

### Datasets and feature extraction

The sample sizes of each of the datasets obtained are listed in table 1. The Original-X and Imbalanced-X datasets have variable number of calls per call type and individual (supplementary table 1).

**Table 1.** Sample sizes of datasets.

| Dataset | Trills | Phees | Food calls |
| --- | --- | --- | --- |
| Original-X | 1246 | 1443 | 4434 |
| Imbalanced-X | 1206 | 1401 | 4419 |
| Balanced-X | 4528 | 3350 | 15372 |
| Balanced197-X | 3152 | 1970 | 3546 |
| Balanced99-X | 1584 | 990 | 1782 |
| Balanced50-X | 800 | 500 | 900 |

We could extract between 3776 to 4553 features from each marmoset call in the Original-X datasets, with 3255, 3395, 3477 features common across trills, phees, and food calls, respectively.

### ML classifiers and feature importance scoring

We monitored the classification loss function of AdaBoost with the addition of every new weak learner. We observed the loss function to plateau at around 500 trees while training AdaBoost to determine sex and around 2500 for other tasks.

#### Balanced vs unbalanced datasets

Classification accuracies ranged from 60.8% to 70.96% for imbalanced datasets versus 71.43% to 83.92% for balanced datasets. Mean ROC-AUCs were higher for all the Balanced-X datasets compared to Imbalanced-X datasets (table 2).

**Table 2.** AdaBoost performance for imbalanced and balanced datasets. Imbalanced-X and Balanced-X datasets were used. Mean ± s.d. are provided for ROC-AUC scores.

| Call | Classes<br>(Individuals) | Imbalanced |  |  | Balanced |  |  |
| --- | --- | --- | --- | --- | --- | --- | --- |
|  |  | Sample size | ROC-AUC (%) | Accuracy (%) | Sample size | ROC-AUC (%) | Accuracy (%) |
| Trills | 16 | 1206 | 92.63 ± 4.30 | 64.84 | 4528 | 98.95 ± 0.91 | 82.91 |
| Phees | 10 | 1498 | 94.36 ± 4.54 | 70.96 | 3550 | 98.55 ± 1.42 | 83.92 |
| Food calls | 18 | 4419 | 92.72 ± 5.08 | 60.8 | 15372 | 97.04 ± 2.31 | 71.43 |

#### Top 20 vs random 20 features

t-SNE plots obtained using top 20 features on the Balanced-X datasets showed visibly better clusters compared to t-SNE plots obtained using random 20 features on the corresponding datasets (figure 2, supplementary tables 2, 3 and 4). The top 20 features provided by AdaBoost gave significantly greater silhouette scores than what would be obtained by chance for trills (Z = 7.0154, p = 1.146e-12), phees (Z = 4.8361, p = 6.621e-7), and food calls (Z = 6.8922, p = 2.747e-12).

**Figure 2.**
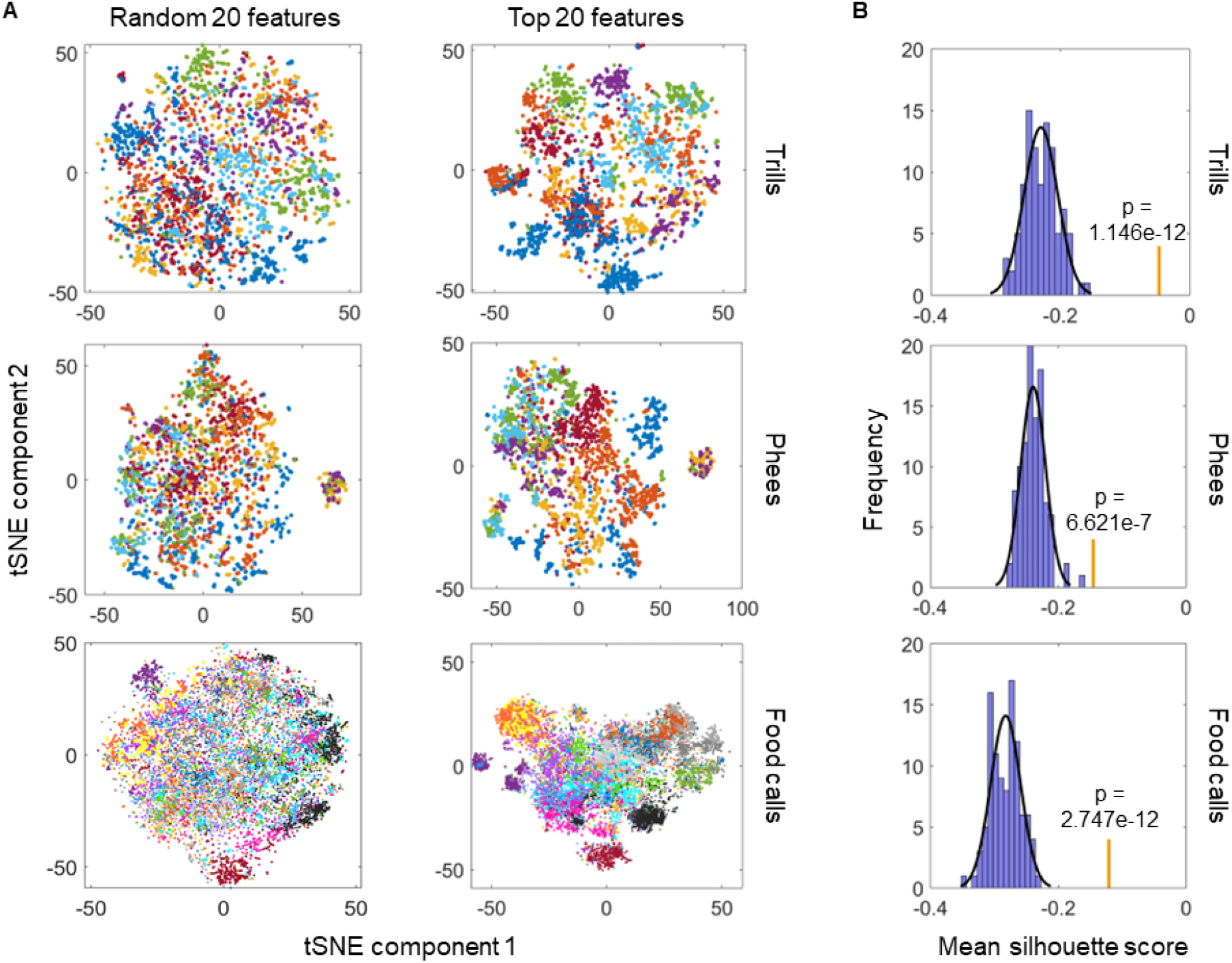
Qualitative and quantitative comparisons of random 20 and top 20 features for clustering data. (A) Qualitative comparison. Figures are t-SNE plots (squared Euclidean distance metric) of Balanced-X datasets using 20 randomly chosen features or the top 20 features for classification by AdaBoost for that call type. Each point is a call, coloured according to the source individual. (B) Quantitative comparison. Blue bars depict frequencies (histogram) of mean silhouette scores obtained after performing t-SNE using 20 random features selected 100 times for that call type (trills/phees/food calls) on Balanced-X datasets. The grey line depicts the Gaussian function fit to the histogram. Orange vertical bars denote the mean silhouette scores obtained after performing t-SNE using top 20 features for classification by AdaBoost for that call type. The p-value shown is the normalised area under the gaussian function to the right of the orange bar.

#### Hierarchical vs non-hierarchical classifiers at large sample sizes

The accuracies of AdaBoost to classify the following datasets based on sex were: Balanced-T = 99.4%, Balanced-P = 96.8%, Balanced-F = 96.7%. The hierarchical classification approach gave significantly better precision and recall scores compared to the non-hierarchical approach for the corresponding Balanced-X datasets (p <0.05 across all call types, Wilcoxon signed-rank test; table 3). A thorough evaluation of the precisions and recalls for each individual by the hierarchical classifier revealed that the trills of 2 individuals - Washington and Wisconsine (twins), had significantly low scores (precisions: 74.23% and 67.46% respectively, recalls: 75.07% and 72.99%, figure 3). The mean precision and recalls for trills for all except these two individuals were as high as 97.79% and 97.07%, respectively. The phee precision and recall scores for Washington and Wisconsine were also slightly lower than that of other females (supplementary figures 1A and 1B). For food calls, the precision and recall scores of these two individuals were similar or higher than the mean scores of all females (supplementary figures 1C and 1D).

**Figure 3.**
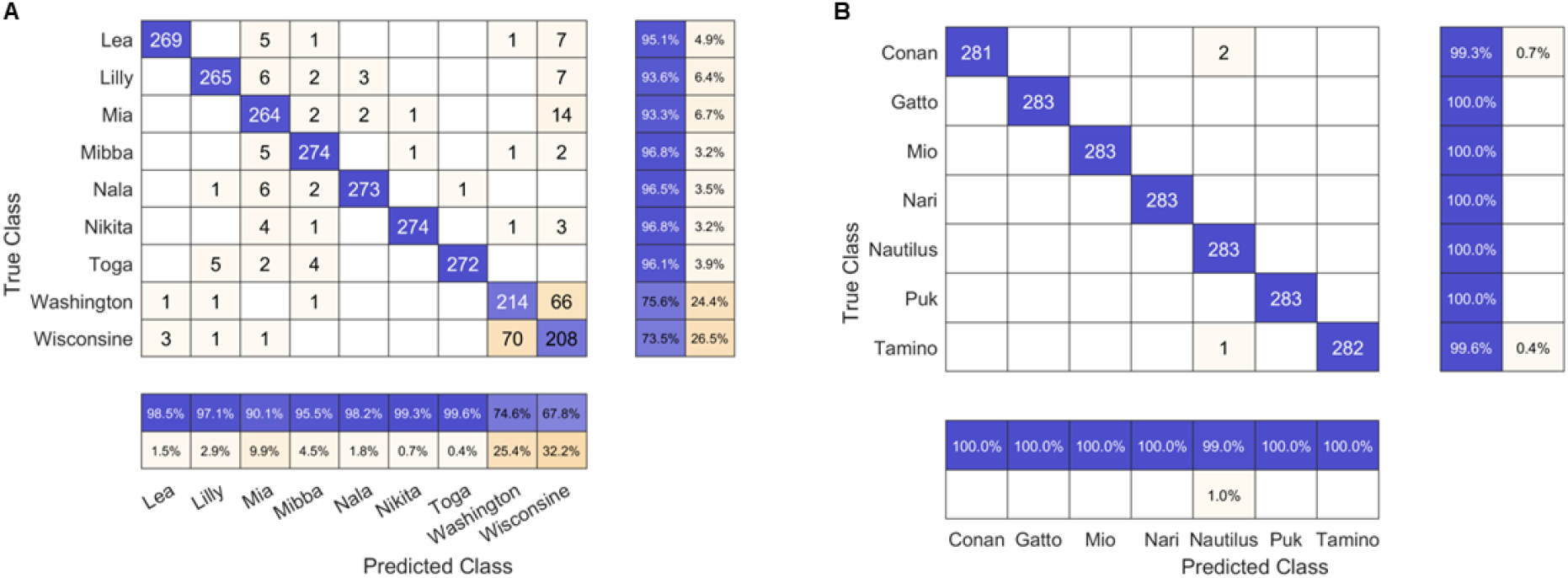
Individual precisions and recalls for determining the source identity from trills by the hierarchical classifier for females (A) and males (B). Confusion matrices are shown, with rows depicting the true source identity and columns representing the prediction made by the hierarchical classifier. The absolute number of calls is shown within the matrices with those correctly classified highlighted in blue and those wrongly classified highlighted in orange (intensity proportional to the number for both). The rows and columns are summarised with the row summary depicting individual precisions in blue and the columns summary showing individual recalls in blue.

**Table 3.** Comparing non-hierarchical and hierarchical approaches for classifying calls based on source identity. Mean ± s.d. precisions and recalls with corresponding p-values for testing the hypotheses: mean precisions/recalls of non-hierarchical = hierarchical.

| Call | Classes (individuals) | Non-hierarchical |  | Hierarchical |  | p-value |  |
| --- | --- | --- | --- | --- | --- | --- | --- |
|  |  | Precision (%) | Recall (%) | Precision (%) | Recall (%) | For precision | For recall |
| **Trills** | 16 | 83.42 ± 11.48 | 82.87 ± 8.63 | 94.42 ± 9.61 | 94.19 ± 8.24 | **0.017** | **0.01** |
| **Phees** | 10 | 84.15 ± 9.84 | 83.93 ± 8.63 | 92.57 ± 3.30 | 92.55 ± 3.79 | **0.027** | **0.037** |
| **Food calls** | 18 | 72.23 ± 15.09 | 71.43 ± 13.13 | 87.21 ± 9.71 | 86.65 ± 5.62 | **0.01** | **0.002** |

#### Hierarchical vs non-hierarchical classifiers at reduced sample sizes

With the decrease in sample size per class, the difference between the mean precisions and recalls of the hierarchical and non-hierarchical approaches reduced for all three call types (figure 4 for precision; see supplementary figure 3 for recall).

**Figure 4.**
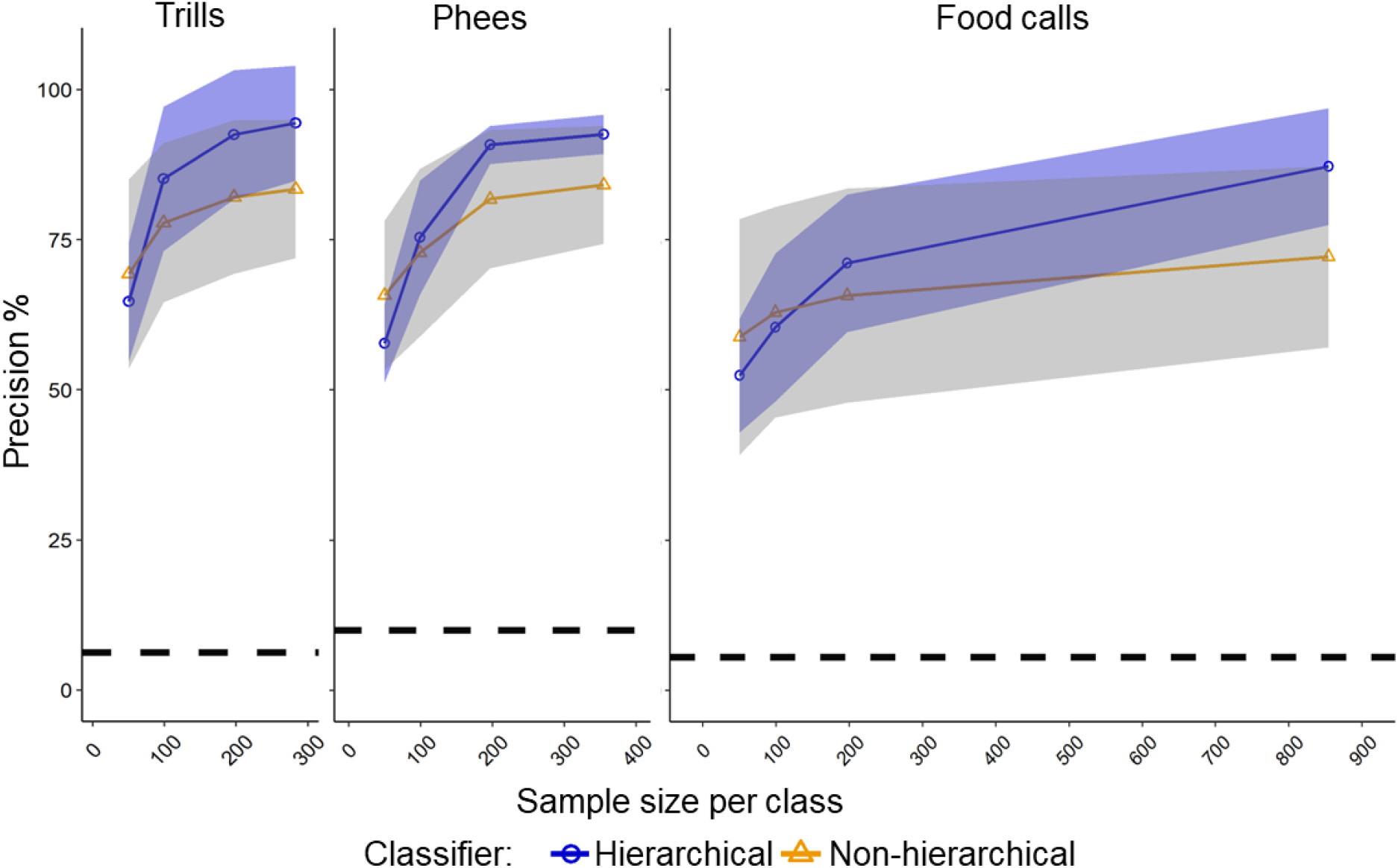
Classifier performance at different sample sizes. Precisions of AdaBoost as a function of sample size per class for trills, phees, and food calls with s.d. represented as shaded regions around lines connecting means. Dashed black lines indicate the chance precision of classification for given call type. Note that the highest sample size per class was obtained by oversampling the minority classes so that they are equal to the majority class (SMOTE, see methods). See supplementary table 1 for sample sizes in our original dataset.

#### Feature selection at various levels of the hierarchical classifier

The features selected by each level of the hierarchy varied with the task. About 30%-60% of the features were utilised solely for determining sex across call types. Only a few features were common for determining individual identities among males and females (figure 5).

**Figure 5.**
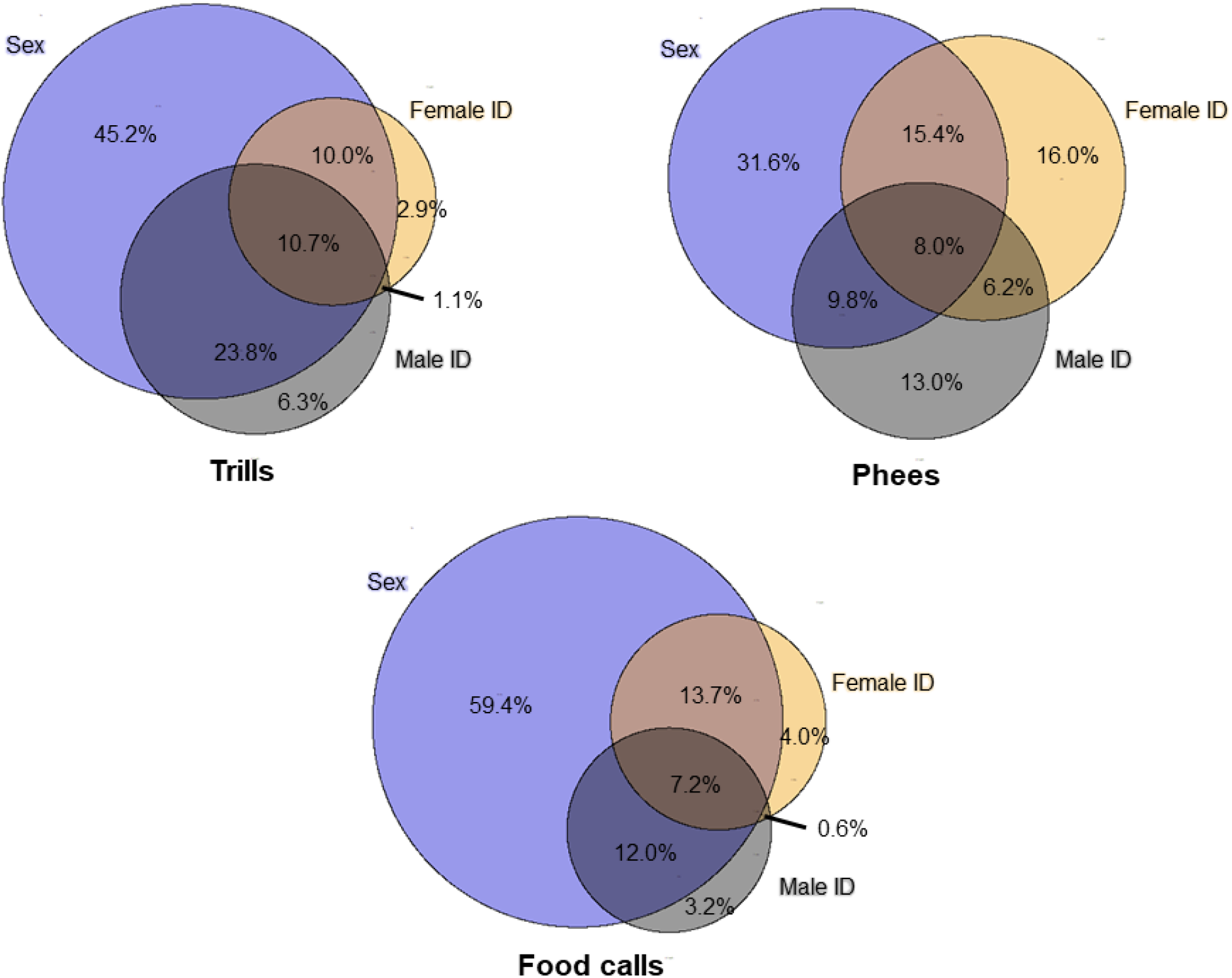
Mutual and distinct features used by hierarchical classifiers at various levels of classification. Venn diagrams denote the set of features used for determining the sex, source identity among females (Female ID), and source identity among males (Male ID). The area of the circle is scaled to the number of features at that level. The percentage of total features used by the hierarchical classifier belonging to each area within the Venn diagram is denoted.

## DISCUSSION

We show that the information about sex encoded in marmoset calls can be used to build hierarchical ML classifiers to determine the source identity with higher precisions and recalls than with their non-hierarchical counterparts. We incorporated such a hierarchical classifier in a pipeline that starts with plentiful feature extraction to provide a rich representation of the calls, followed by meaningful feature selection, and source identification from marmoset vocalisations. Our findings have implications for marmoset communication research and will be beneficial for tracking changes in vocalisations during vocal learning. The same methods can also be extended for efficient source identification in other species.

### Optimising source identification

Of the large number of features extracted with HCTSA, we found that the AdaBoost algorithm reliably selected the most important features that would best cluster the data for its classification task (figure 2, supplementary tables 1, 2, and 3). Intriguingly, the features that were selected varied with the task of the classifier (figure 4). Because all subsequent quantitative analyses of vocalisations depend on feature selection, this step is important for studying animal vocalisations. In contrast to our approach, more traditional approaches typically use a standard set of acoustic features across all tasks e.g. [62,63] or [64,65]. This may lead to a suboptimal representation of the call for a specific task at hand, and thus lower classification accuracy. Hence, despite dealing with the same call type, a customised set of features seems beneficial for optimal performance in a specific classification task. Secker et al. [66] have previously used a hierarchical classifier with independent feature selection by the components to classify proteins successfully. They suggest that independent feature selection helps maintain high predictive performance while improving computational efficiency. In our case, the independent feature selection arguably enabled the hierarchical classifier to efficiently utilise broader-category cues (i.e., sex) for classification, hence boosting performance over its non-hierarchical counterpart. AdaBoost thus provides task specific and flexible feature extraction with a customised set of features for every dataset and can be applied to a wide range of animal vocalisation datasets. However, many of these features are complex, with some only defined by the mathematical operations performed on the audio waveform for obtaining the feature, hence less interpretable. Therefore, feature interpretability is a limitation. However, this is compensated by decision tree based AdaBoost, making the interpretability of our approach better than many “black-box” models such as deep neural networks [67].

Even with the substantial number of features provided by HCTSA for AdaBoost to work with, AdaBoost performed poorly on the unbalanced dataset with accuracies below 70% (table 2). A large body of ML literature points out to the problem of class-unbalanced datasets and its solutions [68,69]. Recent studies and reviews have emphasised the use of data augmentation and balancing techniques to improve ML accuracy when handling acoustic data [70–72]. Consistent with other studies [73–75], balancing the datasets significantly improved the performance of AdaBoost across call types. Therefore, data balancing using tools like SMOTE combined with random undersampling is an important step before running MLAs on any dataset.

Classifying data points from a noisy dataset to a large number of classes (10 – 18 different individuals in our case) is often a demanding task for an MLA. Here, we broke the problem of classifying calls to over ten sources into a first classification problem of assigning the sex of the source, and only then, given the sex, classifying source identity in a second step (figure 1). We showed that at large sample sizes, mean precisions and recalls across datasets increased by more than eight percentage points with this hierarchical approach (table 3). We found the accuracies of the hierarchical classifier on the largest balanced datasets to remain satisfactory and higher than most recent studies classifying animal vocalisations using MLAs [62,76,77], even though the number of samples per class of data was lower than those studies. The same was reflected in the performance of the hierarchical classifier on the originally collected calls (supplementary figure 2).

The difference in precisions and recalls between the hierarchical and non-hierarchical approaches diminished as the sample sizes decreased suggesting that the hierarchical classifier requires exposure to enough data to perform significantly better than its non-hierarchical counterpart (figure 4, supplementary figure 3). Lower sample sizes pose a higher risk of error cascading in the hierarchical classifier, and this has been identified in similar classifiers developed for text classification [78,79]. This is due to the error at the top of the hierarchy propagating down the levels of the hierarchical classifier. However, this limitation only arises at the level of the training dataset.

The requirement of a large training dataset and the presence of a small collected dataset can almost always be bridged. The median sample sizes per class in our Imbalanced-X datasets were 47 for trills, 83 for phees and 173 for food calls (supplementary table 1). Using this small, highly imbalanced dataset, we could generate larger balanced datasets and train and test our classifiers on them (see methods). Therefore, to obtain optimal performance from the hierarchical classifier, one need not necessarily require a large sample size to begin with (see performance of the hierarchical classifier on the Imbalanced-X dataset in supplementary figure 2).

### Implications for understanding the marmoset communication system

The classification precisions and recalls for food calls were low compared to trills and phees despite over three times higher sample sizes (table 3). In particular, food calls required a higher sample size per class value to be classified as accurately as trills and phees (figure 4). On the one hand, food calls are likely a highly heterogeneous group of call types [24]. Moreover, they are predominantly produced in bouts of multiple repeated call units [21]. Furthermore, each call unit is much shorter than a trill or a phee [21], therefore providing reduced information for time series analysis. The poor classification results could thus arise because some of the source identity information may well be encoded at the level of the bout. This is testable in the future by repeating the classification procedure on bouts of food calls.

Intriguingly, the mean precision and recall for determining source identity from trills when considering all except two individuals was as high as 98.38% and 97.65%, respectively (figure 3). The two marmosets with low scores happened to be twins of the same sex. Thus, the low classification scores were likely due to the high vocal similarity between the calls of the twins. The same pattern was present to a lesser extent for phee calls but not for food calls (supplementary figure 1). Whereas the food calls may require further scrutiny at different levels of analyses (see above), the contrast between trills and phees is interpretable with regard to their biological function. Signalling identity is essential for the long-distance phee calls that individuals typically use to establish acoustic contact when visual contact is not possible [80]. In contrast, trill calls are given in close proximity to a social partner, and the caller’s identity is thus redundant because the marmosets may also use visual or olfactory cues to recognise the partner [81]. It may thus well be that the twins actively diverged from each other in their phee calls but not in their trill calls. This is consistent with a recent study [50] on newly paired marmosets that found that partners would converge in the structure of their phee calls. However, newly formed pairs that had initially very similar phee calls before pairing diverged rather than converged in their call structure, supposedly to make themselves better distinguishable. In the future, our approach will be beneficial for precisely tracking changes in vocalisations in such situations, as well as during ontogeny because it would be possible to track changes in features that most differentiate two individuals. More precise tracking of vocal changes will enable us to identify why such a phenomenon might have gotten selected, both in adults and immatures [82,83]. Getting to the bottom of the factors facilitating vocal learning and other enhanced communicative capabilities of cooperatively breeding primates will better inform us about the evolutionary origins of language [23].

Finally, we know that callitrichids can differentiate between vocalisations originating from cage-mates versus foreign individuals and from males versus females [48,49]. An intriguing question is how they achieve that, and whether their decision-making process may likewise be hierarchically structured with broader-category cues used as a first distinction. Multiple studies on humans allude to the hierarchical nature of decision-making in various contexts [84–89]. Some frog species seem to employ hierarchical decision-making for prey capture [90], a hierarchical decision-making model appears to best explain the strategies used by two rhesus macaques while playing a slightly modified, semi-controlled adaptation of the video game ‘Pacman’ [91], a hierarchy of multimodal cues are used by sea lions during mother-offspring recognition [92,93], and evidence from fruit flies, locusts, and zebrafish suggest that they break down a complex problem of deciding between spatially distributed options into a series of smaller problems [94]. Hierarchical decision-making may thus be a widespread feature of cognitive processing. Given that sex could be attributed with extremely high precision by our classifier, we hypothesise that marmosets, too, use these sex-based cues for efficient source identification from calls, and they are doing so in a hierarchical manner, similar to the hierarchical classifier. Cognitive and psychological experiments will be required to test this hypothesis.

Overall, we demonstrated the capabilities of our pipeline for determining source identity from marmoset vocalisations by breaking down the larger classification problem into a hierarchy of smaller, easier problems. Although our hierarchical approach focused on sex-based cues in marmoset vocalisations, the idea is for it to be generalisable to all broad-category cues that animal vocalisations may provide. In the future, we hope to extend the idea to larger, more complex animal vocalisation datasets that would require multiple levels in the hierarchy and would utilise other cues, such as the age and social status of the animal, for efficient source identification. Finally, the ability to select customised features for every dataset and task, combined with the supervised nature of learning, makes our pipeline highly flexible and extendable for analysing vocalisations of diverse animal species.

## ACKNOWLEDGEMENTS AND FUNDING

This work was supported by the Swiss National Science Foundation (grant number 31003A_149796, the NCCR Evolving Language, agreement number 51NF40_180888), and the European Research Council (ERC) under the European Union’s Horizon 2020 research and innovation programme (grant agreement No 101001295). NP was a recipient of a grant by the A. H. Schultz foundation.

