## Supplementary materials for "Optimising source identification from marmoset vocalisations with hierarchical machine learning classifiers"

**Supplementary table 1. The Original-X dataset.** Values denote the number of calls per call type (column) and individual (row). The calls discarded from the Original-X dataset to obtain the Imbalanced-X dataset are shown in red.

| Sex | Individual | Trills | Phees | Food calls |
|---|---|---|---|---|
| Female | Jaja | 0 | 0 | 2 |
| | Lea | 47 | 1 | 36 |
| | Lilly | 55 | 62 | 403 |
| | Mia | 43 | 7 | 78 |
| | Mibba | 47 | 66 | 545 |
| | Nala | 37 | 0 | 99 |
| | Nikita | 62 | 16 | 286 |
| | Toga | 78 | 8 | 164 |
| | Washington | 243 | 70 | 111 |
| | Wisconsine | 283 | 335 | 182 |
| Male | Conan | 85 | 355 | 203 |
| | Craken | 20 | 96 | 287 |
| | Gatto | 52 | 171 | 509 |
| | Kyros | 18 | 11 | 64 |
| | Membo | 2 | 50 | 121 |
| | Mio | 34 | 4 | 98 |
| | Nari | 38 | 10 | 333 |
| | Nautilus | 36 | 128 | 46 |
| | Puk | 25 | 12 | 13 |
| | Tamino | 42 | 41 | 854 |
| **Total (Original-X)** | | **1246** | **1443** | **4434** |
| **Total analysed (Imbalanced-X)** | | **1206** | **1401** | **4419** |

**Supplementary table 2. HCTSA functions for extracting top 20 features from trills.** Ranks and descriptions of HCTSA functions used to extract top 20 features from trills. Descriptions taken from Fulcher and Jones (2017).

| Feature rank | HCTSA function | Description |
|---|---|---|
| 1 | MF-arfit-1-8-sbc-sumA | Autoregressive model fitting. Optimum order is chosen from the range 1 to 8. Detects inherent repeating patterns. |
| 2 | SY-SlidingWindow-mom3-ent10-10 | Time series cut into 10 equal windows. Skewness calculated for each window. Entropy of skewness values given as output. |
| 3 | SY-RangeEvolve-nuql1000 | Describes how the range of the time series (i.e range of pressure values in the acoustic waveform) changes with time. |
| 4 | EX-MovingThreshold-1-002-stdkickf | Tracks the number of extreme events using a dynamic 'moving threshold'. Every new point in the timeseries is classified as either 'extreme' or 'not extreme' based on the history of extreme events. The standard deviation of this threshold across the timeseries is given as output |
| 5 | EX-MovingThreshold-01-002-mediankickf | Tracks the number of extreme events using a dynamic 'moving threshold'. Every new point in the timeseries is classified as either 'extreme' or 'not extreme' based on the history of extreme events. The median of this threshold across the timeseries is given as output. |
| 6 | MF-arfit-1-8-sbc-A4 | Autoregressive model fitting. Optimum order is chosen from the range 1 to 8. Detects inherent repeating patterns. |
| 7 | SY-SpreadRandomLocal-200-100-stdskew | 100 segments each 200ms long are randomly selected, skewness calculated, and the standard deviation of the skewness values given as output. |
| 8 | SB-MotifThree-diffquant-ccaa | Performs coarse graining of the time series. |
| 9 | SY-LocalGlobal-std-p5 | Compares the standard deviation of the first 1/5th of the time series to that of the entire time series |
| 10 | PH-Walker-momentum-2-sw-taudiff | A hypothetical particle that moves along the length of the time series is simulated. The movement is dynamic and depends on the pressure value of the acoustic waveform at that time. The trajectory of the particle is summarised and compared to that of the actual timeseries. |

| | | |
|---|---|---|
| 11 | IN-AutoMutualInfoStats-40-gaussian-pcrossmedian | Median value of the auto-mutual information |
| 12 | SC-MMA-0--n5-5-maxHurstExponent | Multifractal scaling of the time series performed. Maximum Hurst exponent given as output |
| 13 | TSTL-localdensity-5-40-ac-2-minden | Provides estimates of local densities while performing time-delay embedding |
| 14 | NL-TSTL-ReturnTime-10-1-1-n1-ac-8-iqr | Time taken for the pressure value of the acoustic waveform to return to the same location. Provides evidence for periodicities in the data. |
| 15 | SY-SpreadRandomLocal-100-100-stdsampen1-015 | 100 segments each 100ms long are randomly selected, sample entropy calculated, and standard deviation of those values returned. |
| 16 | SC-MMA-0--n5-5-qHurstTrend | Multifractal scaling of the time series performed. $5^{th}$ quantile of Hurst exponent given as output |
| 17 | CO-TranslateShape-rectangle-2-pts-ones | A rectangle is moved along the length of the time series and statistics performed on all points lying inside the rectangle. |
| 18 | CO-TranslateShape-rectangle-2-pts-fives | A rectangle is moved along the length of the time series and statistics performed on all points lying inside the rectangle. |
| 19 | SC-MMA-0--n5-5-meanHurstExponent | Multifractal scaling of the time series performed. Mean Hurst exponent given as output |
| 20 | SY-SpreadRandomLocal-200-100-stdac2 | 100 segments each 200ms long are randomly selected, autocorrelation coefficient calculated, and the standard deviation of the autocorrelation coefficients values given as output. |

**Supplementary table 3. HCTSA functions for extracting top 20 features from phees.** Ranks and descriptions of HCTSA functions used to extract top 20 features from phees. Descriptions taken from Fulcher and Jones (2017).

| Feature rank | HCTSA function | Description |
|---|---|---|
| 1 | SY-SpreadRandomLocal-200-100-stdskew | 100 segments each 200ms long are randomly selected, skewness calculated, and the standard deviation of the skewness values given as output. |
| 2 | SY-SlidingWindow-mom3-ent10-10 | Time series cut into 10 equal windows. Skewness calculated for each window. Entropy of skewness values given as output. |
| 3 | NL-crptool-fnn-10-2-1-firstunder01 | Inspects the false nearest neighbour function using time-delayed embedding. |
| 4 | IN-AutoMutualInfoStats-40-kraskov1-4-amiac1 | Autocorrelation of auto-mutual information values obtained using the Kraskov estimator. |
| 5 | SY-SpreadRandomLocal-200-100-meanac1 | 100 segments each 200ms long are randomly selected, autocorrelation coefficient calculated, and the mean of the autocorrelation coefficients given as output. |
| 6 | IN-AutoMutualInfoStats-40-gaussian-amiac1 | Autocorrelation of auto-mutual information values obtained using the Gaussian estimator. |
| 7 | IN-AutoMutualInfoStats-40-kraskov1-4-ami7 | Auto-mutual information calculated using the Kraskov estimator. |
| 8 | EX-MovingThreshold-01-01-meankickf | Tracks the number of extreme events using a dynamic ‘moving threshold’. Every new point in the timeseries is classified as either ‘extreme’ or ‘not extreme’ based on the history of extreme events. The mean of this threshold across the timeseries is given as output. |
| 9 | length | Length of the time series |
| 10 | IN-AutoMutualInfoStats-40-kraskov1-4-ami8 | The autocorrelation of the auto-mutual information function of the time series |
| 11 | SC-MMA-0--n5-5-maxHurstExponent | Multifractal scaling of the time series performed. Maximum Hurst exponent given as output |
| 12 | MF-arfit-1-8-sbc-aerr-mean | Fits an autoregressive model of an optimal order |

| | | |
|---|---|---|
| 13 | SY-SlidingWindow-mom3-ent10-2 | Time series cut into 10 equal windows. Skewness calculated for each window. Entropy of skewness values given as output. |
| 14 | NL-crptool-fnn-10-2-1-fnn8 | Inspects the false nearest neighbour function using time-delayed embedding. |
| 15 | SY-SpreadRandomLocal-100-100-stdskew | 100 segments each 100ms long are randomly selected, skewness calculated, and the standard deviation of the skewness values given as output. |
| 16 | SY-RangeEvolve-totnuq | Describes how the range of the time series (i.e range of pressure values in the acoustic waveform) changes with time. |
| 17 | NL-TSTL-dimensions-50-ac-fnnmar-bc-minmmax | Correlation dimension of a time-delay embedded time series. |
| 18 | MF-arfit-1-8-sbc-meanA | Autoregressive model fitting. Optimum order is chosen from the range 1 to 8. Detects inherent repeating patterns. |
| 19 | NW-VisibilityGraph-norm-entropy | Constructs a visibility graph of the time series and returns the entropy of the resulting network |
| 20 | PH-ForcePotential-dblwell-1-02-01-ac10 | Taken from the particle in a double well concept. Forces experienced are a function of the pressure values in the acoustic waveform. |

**Supplementary table 4. HCTSA functions for extracting top 20 features from food calls.** Ranks and descriptions of HCTSA functions used to extract top 20 features from food calls. Descriptions taken from Fulcher and Jones (2017).

| Feature rank | HCTSA function | Description |
|---|---|---|
| 1 | SC-MMA-0--n5-5-meanHurstExponent | Multifractal scaling of the time series performed. Mean Hurst exponent given as output. |
| 2 | PH-Walker-runningvar-15-50-sw-taudiff | A hypothetical particle that moves along the length of the time series is simulated. The movement is dynamic and depends on the pressure value of the acoustic waveform at that time. The trajectory of the particle is summarised and compared to that of the actual timeseries. |
| 3 | SC-MMA-0--n5-5-maxHurstExponent | Multifractal scaling of the time series performed. Maximum Hurst exponent given as output. |
| 4 | mean | Mean (pressure) value of the waveform |
| 5 | NL-TSTL-ReturnTime-005-1-005-n1-1-3-std | Time taken for the pressure value of the acoustic waveform to return to the same location. Provides evidence for periodicities in the data. |
| 6 | SC-MMA-0--n5-5-stdHurstExponent | Multifractal scaling of the time series performed. Standard deviation of Hurst exponent given as output. |
| 7 | NL-TSTL-GPCorrSum2-n1-05-100-20-ac-fnnmar-robfit-sea2 | Correlation sum scaling by Grassberger-Proccacia algorithm |
| 8 | SC-MMA-0--n5-5-qHurstTrend | Multifractal scaling of the time series performed. 5$^{th}$ quantile of the Hurst exponent given as output. |
| 9 | MF-arfit-1-8-sbc-A4 | Autoregressive model fitting. Optimum order is chosen from the range 1 to 8. Detects inherent repeating patterns. |
| 10 | CP-ML-StepDetect-l1pwc-005-E | Analysis of discrete steps in a time series. Gives information about discrete steps in the signal. |
| 11 | DN-ProportionValues-zeros | Returns statistics on the values of the data vector: the proportion of zeros, the proportion of positive values, and the proportion of values greater than or equal to zero. |

| | | |
|---|---|---|
| 12 | NL-TSTL-GPCorrSum2-n1-05-40-20-ac-fnnmar-robfit-sea2 | Correlation sum scaling by Grassberger-Proccacia algorithm |
| 13 | SY-RangeEvolve-nuql1000 | Describes how the range of the time series (i.e range of pressure values in the acoustic waveform) changes with time. |
| 14 | SB-BinaryStats-mean-longstretch0 | Provides information about the coarse-grained behavior of the time series using binary symbolisation to 0s and 1s. |
| 15 | NW-VisibilityGraph-horiz-expnlogL | Constructs a visibility graph of the time series and returns various statistics on the properties of the resulting network. |
| 16 | NL-TSTL-ReturnTime-10-1-1-n1-ac-8-rangecgdist | Time taken for the pressure value of the acoustic waveform to return to the same location. Provides evidence for periodicities in the data. |
| 17 | NL-crptool-fnn-10-2-ac-firstunder02 | Inspects the false nearest neighbour function using time-delayed embedding. |
| 18 | NL-TSTL-ReturnTime-10-1-1-n1-ac-8-std | Time taken for the pressure value of the acoustic waveform to return to the same location. Provides evidence for periodicities in the data. |
| 19 | SY-SlidingWindow-mom3-ent10-10 | Time series cut into 10 equal windows. Skewness calculated for each window. Entropy of skewness values given as output. |
| 20 | MF-arfit-1-8-sbc-minper | Autoregressive model fitting. Optimum order is chosen from the range 1 to 8. Detects inherent repeating patterns. |

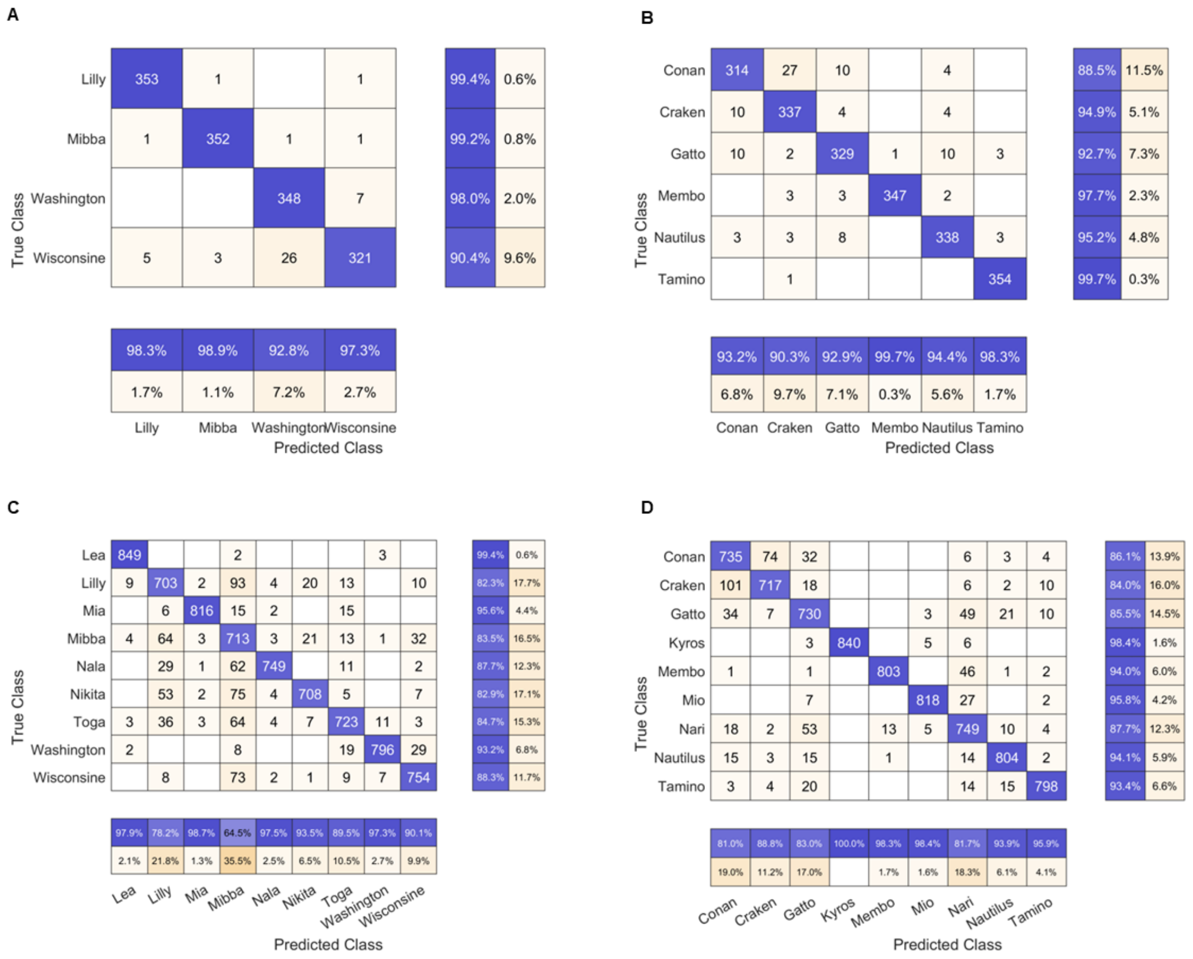

**Supplementary figure 1. Individual precisions and recalls for determining the source identity from phees (A, B) and food calls (C, D) by the hierarchical classifier for females (A, C) and males (B, D).** Confusion matrices are shown, with rows depicting the true source identity and the columns depicting the prediction made by the hierarchical classifier. The absolute number of calls are shown within the matrices with those correctly classified highlighted in blue and those wrongly classified highlighted in orange (saturation proportional to the number for both). The rows and columns are summarised with the row summary depicting individual precisions in blue and the columns summary depicting individual recalls in blue.

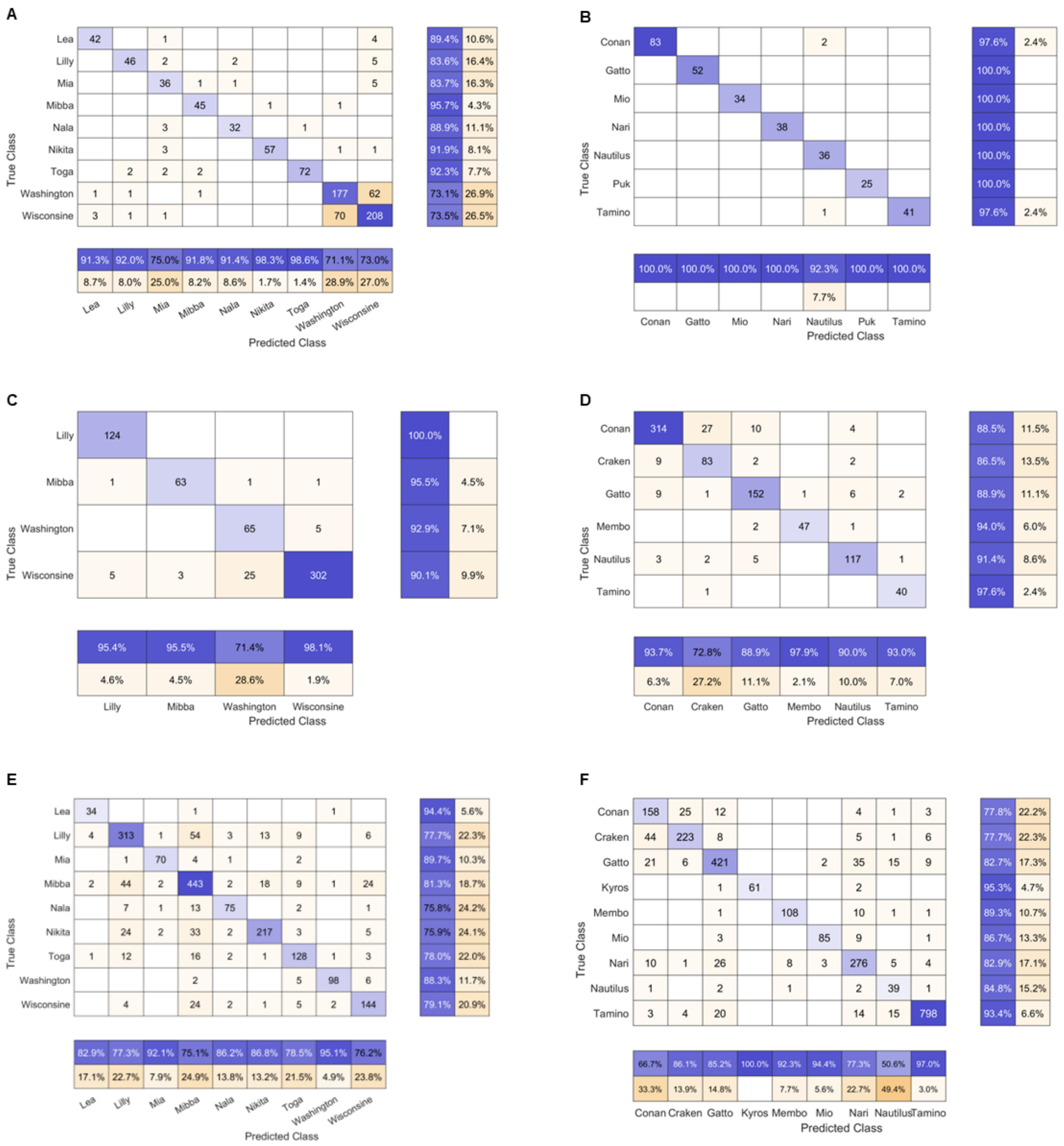


**Supplementary figure 2. Individual precisions and recalls for determining source identity on the originally collected trills (A, B), phees (C, D) and food calls (E, F) (see supplementary table 1), by the hierarchical classifier for females (A, C, E) and males (B, D, F).** Confusion matrices are shown, with rows depicting the true source identity and the columns depicting the prediction made by the hierarchical classifier. The absolute number of calls are shown within the matrices with those correctly classified highlighted in blue and those wrongly classified highlighted in orange (saturation proportional to the number for both). The rows and columns are summarised with the row summary depicting individual precisions in blue and the columns summary depicting individual recalls in blue.

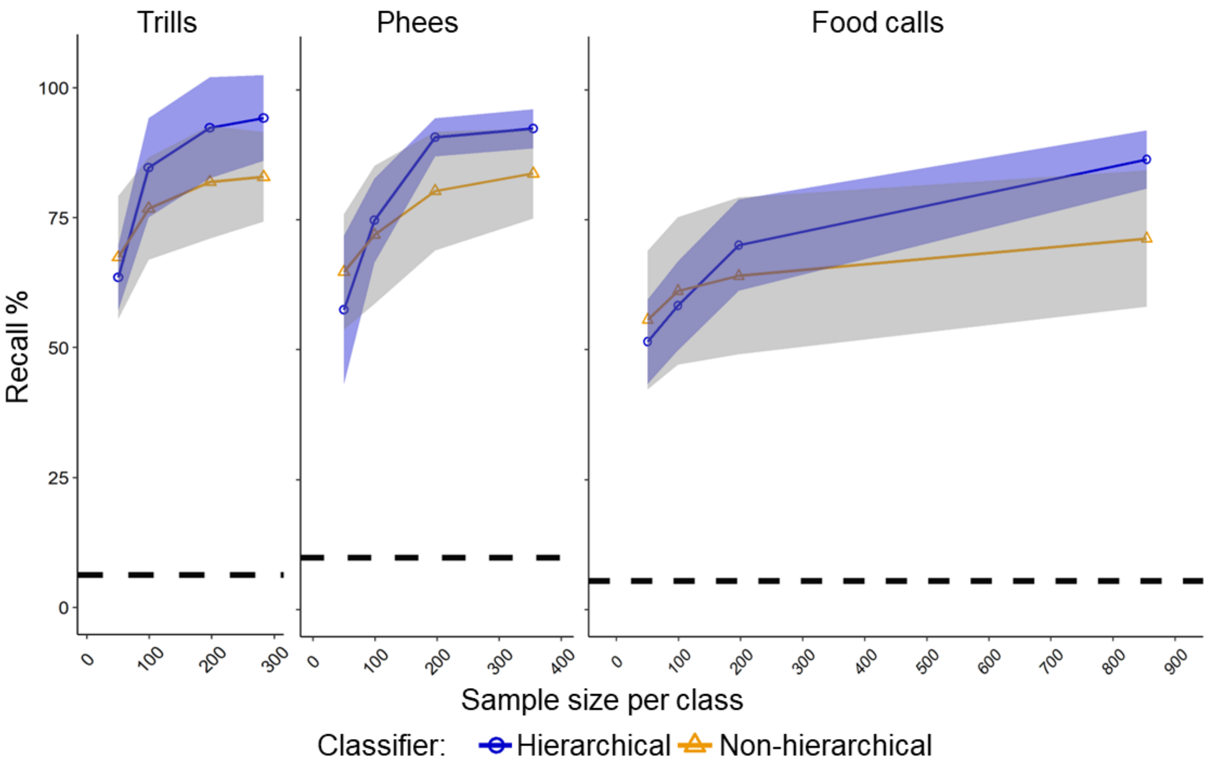


**Supplementary figure 3. Classifier performance at different sample sizes.** Recalls of AdaBoost as a function of sample size per class for trills, phees, and food calls with s.d. represented as shaded regions around lines connecting means. Dashed black lines indicate the chance precision of classification for a given call type.
